# Myocardial infarction injury is exacerbated by nicotine in vape aerosol exposure

**DOI:** 10.1101/2024.07.29.605355

**Authors:** Clarissa Savko, Carolina Esquer, Claudia Molinaro, Sophie Rokaw, Abraham Grant Shain, Faid Jaafar, Morgan K. Wright, Joy A. Phillips, Tyler Hopkins, Sama Mikhail, Abigail Rieder, Ariana Mardani, Barbara Bailey, Mark A. Sussman

## Abstract

**Rationale:** Vaping is touted as a safer alternative to traditional cigarette smoking but the full spectrum of harm reduction versus comparable risk remains unresolved. Elevated bioavailability of nicotine in vape aerosol together with known risks of nicotine exposure may result in previously uncharacterized cardiovascular consequences of vaping.

**Objective:** Assess the impact of nicotine exposure via vape aerosol inhalation upon myocardial response to infarction injury.

**Methods and Results:** Flavored vape juice containing nicotine (5 mg / ml) or vehicle alone (0 mg) was delivered using identical 4-week treatment protocols. Mice were subjected to acute myocardial infarction injury and evaluated for outcomes of cardiac structure and function. Findings reveal that nicotine exposure leads to worse outcomes with respect to contractile performance regardless of sex. Non-myocyte interstitial cell accumulation following infarction significantly increased with exposure to vape aerosol alone, but a comparable increase was not present when nicotine was included.

**Conclusions:** Myocardial function after infarction is significantly decreased after exposure to nicotine vape aerosol irrespective of sex. Comparable loss of contractile function was not observed in mice exposed to vape aerosol alone, highlighting the essential role of nicotine in loss of contractile function. Increased vimentin immunoreactivity was observed in the vape alone group compared to control and vape nicotine. The correlation between vaping, interstitial cell responses, and cardiac remodeling leading to impaired contractility warrants further investigation. Public health experts seeking to reduce vaping-related health risks should consider messaging that highlights the increased cardiovascular risk especially with nicotine-containing aerosols.

## Introduction

The undeniable health consequences for smoking traditional cigarettes drove development of electronic cigarettes (e-cigarettes) intended to provide smokers with a “harm reduction” alternative electronic nicotine delivery system (ENDS) device. This laudable goal to benefit current smokers seeking a healthier lifestyle was unfortunately counterbalanced by adoption of these devices by “never smokers” including adolescents. A recently completed survey of middle and high school students in the U.S.A. found 10% e-cigarette use with the vast majority consuming flavored e-cigarettes at 89.4%^1^. In comparison, American adult e-cigarette usage decreases with age at approximately 10%, 5%, and 3% for 18-29, 30-44, and 45-59 years, respectively.^2^ The vast majority of adult users are either previous smokers or engage in dual use of both conventional and e-cigarettes. From these surveys it is apparent that e-cigarettes are prevalent in American society and continued studies are warranted to delineate health benefits for smokers to switch as well as the risks for those who adopt e-cigarettes as a lifestyle choice. While e-cigs are often thought to be a “harm reduction” alternative to help smokers abstain from combustible tobacco cigarettes, these ENDS devices are not without health risks. Furthermore, the ability to manipulate nicotine concentration in vape juice to higher levels than conventional cigarettes as well as higher bioavailability of chemically modified nicotine warrants additional concerns of stronger addiction and increased physiological consequences.

Adverse impacts of vaping upon the cardiopulmonary system have been demonstrated in multiple experimental studies^3-8^ that mirror clinical case reports from heavy vape users in several important ways.^3,9-19^ Notably, cardiovascular structure and function can deteriorate along with pathological changes in the lungs, including inflammation associated with exposure to vape juice flavorings and additives. The highly variable composition of vape aerosols coupled with lack of standardized vaping protocols have contributed to variability in reported outcomes.^20-23^ Despite inherent challenges to model human vaping behavior in experimental models, consensus is emerging that vaping presents a potential health risk, although the full extent and specific consequences are not easily predictable and depend upon a plethora of individual-specific factors.

The full spectrum of physiological responses to individual vape aerosol constituents continues to be revealed by ongoing investigation, but nicotine has been studied for decades with extensive reviews of cumulative literature.^24-29^ Many studies were conducted in the context of smoking tobacco, rendering conclusions on the impact of nicotine impossible to completely dissociate from the multitude of other chemical compounds in cigarette smoke. Nevertheless, nicotine alone exerts very profound and specific actions, including alteration of wound healing and inflammation^30-32^, fibrosis with collagen deposition^33-35^, physiological effects on the cardiovascular system^24,36,37^, as well as cellular and molecular changes in cardiovascular and pulmonary tissues.^37-41^ Multifaceted actions of nicotine *in vivo* coupled with ENDS devices warrants scrutiny to assess disruption of normal physiologic responses.

Current myocardial biology dogma asserts that the inflammatory response to ischemic injury is a critical determinant of tissue remodeling involving fibrosis and scarring.^42-44^ Recent studies point to post-infarction myocardial healing modulation by the inflammatory response to injury, particularly in early phases of mounting the fibrotic response.^45^ The immunomodulatory action of nicotine may impair healing following tissue injury^46,47^ with concomitant pro-fibrotic action^33-35^. Indeed, there is evidence supporting worsened outcomes in cardiac structure and function resulting from smoking or nicotine exposure prior to infarction^26,48-52^, but specific contribution of nicotine in exacerbating myocardial injury is poorly defined. This issue is particularly germane to vape aerosol exposure where bioavailability of nicotine can far exceed that of traditional smoking.^53-55^ Thus, pre-existing exposure to nicotine via vape aerosol inhalation could predispose the heart to increased damage following subsequent pathological stress such as ischemic injury. An altered inflammatory response due to long term systemic exposure to circulating vape aerosol products could compromise myocardial capacity to withstand supervening acute injury such as infarction.

The resultant outcome of any injury response is influenced by many factors, and the complex interplay of nicotine and vape aerosol constituents to disrupt normal homeostatic mechanisms of tissue repair has received scant attention particularly in the myocardial context. This study was undertaken to investigate if prior vape aerosol exposure worsens the outcome of acute cardiac ischemic injury with respect to cardiac structure, function, and fundamental injury responses of tissue fibrosis and inflammation.

## Methods

### Study animals

Four-week-old male and female C57BL/6J mice were purchased from (Jackson Laboratory, catalog #00064) and housed 4 mice per static cage. Ambient temperature was 70– 72 °F, on a 12-h light dark cycle with automatic light control. Mice were supplied with Rodent Maintenance Diet (Teklad Global 14% Protein, Madison WI) and water ad libitum. Animal protocols and experimental procedures were approved by the Institutional Animal Care and Use Committee at San Diego State University.

### Mouse vaping inhalation protocol

Six-to eight-week-old mice were exposed to 70/30 ratio of propylene glycol (PG) and vegetable glycerin (VG) with 5 mg/mL of nicotine salts, watermelon flavor, and menthol prepared as directed by the manufacturer (Wizard Labs, San Diego, CA). The vape juice was inserted into pods in JUUL pens in whole body exposure chambers (inExpose; SCIREQ, Montreal, Canada). Mice were exposed to 3 second puffs every 20 seconds at 1.8 L/minute, intake rate. Exhaust pumps for fresh air flow rate was 2.5 and 1.5 L/minute for the 4-hour duration of the vaping profile based upon human vaping topography recommended by Farsalinos et al. concluding “Four-second puffs with 20–30 s interpuff interval should be used when assessing electronic cigarette effects in laboratory experiments, provided that the equipment used does not get overheated.”^21^ In addition, puff duration parameters and frequency are within the reported range of human vaping topography from real time characterization of electronic cigarette use in the 1 Million Puffs Study.^56^ Animals were exposed for 4 hours/day, 5 days/week for 4 weeks. Cotinine levels were elevated in both males and females but not significantly different at 62.48 ±53.76 versus 71.20 ±39.14, respectively using (Supplemental Figure 1). Animal protocols and experimental procedures were approved by the Institutional Animal Care and Use Committee at San Diego State University.

### Infarction Protocol

Myocardial infarctions were carried under 2% isoflurane (Victor Medical, catalog #NDC 57319-474-06). The 3rd and 4th ribs were separated enough to get adequate exposure of the operating region, but the ribs were kept intact. The heart was squeezed out by pressing the thorax lightly and the left anterior descending artery (LAD) was ligated at the distal diagonal branch with a 7–0 suture. Infarction was confirmed by blanching of anterior left myocardium wall. The hearts were immediately placed back into the intrathoracic space followed by muscle and skin closure. Animals in sham group (*n* = 12) received a comparable surgical procedure without LAD ligation or injection. Each animal received 20 µL analgesic treatment with 0.3 mg/mL Buprenex (Victor Medical, catalog #12496-0757) at time of surgery and 12 h post-surgery.

### Echocardiography

Transthoracic echocardiography was performed on lightly anesthetized mice under isoflurane (1.0–2.0%, Abbot Laboratories) using a Vevo 2100 (VisualSonics). Hearts were imaged in the 2D parasternal long-axis (PSLAX) view, and M-mode echocardiography of the mid-ventricle was recorded at the level of papillary muscles to calculate fractional shortening (FS). From the recorded M-mode images the following parameters were measured: ejection fraction (EF), left ventricular (LV), anterior wall thickness (AWT), LV posterior wall thickness (PWT), LV internal diameter (LVID), and LV volume in diastole (index: d) and systole (index: s).

### Cardiac histology

Following anesthetization of the mice by Ketamine, hearts were arrested in diastole and perfused with formalin for 15 min at 80–100 mmHg via retrograde cannulation of abdominal aorta. Retroperfused hearts were removed from the thoracic cavity and fixed overnight in formalin at room temperature. The hearts were processed for paraffin embedding and sectioned in the coronal orientation at 5 µm thickness at room temperature. The heart sections were stained with Harris Hematoxylin and Eosin Phloxine to visualize morphometric and structural changes. Images were obtained by a Keyence BZ-x600 microscope using XY stage tile scan and stitched using Keyence Analysis software and subsequently analyzed using ImageJ software.

### Infarct measurement

Infarcted hearts sectioned in the coronal orientation were stained with H&E and tile scans were acquired with the Keyence BZ-x600. Percent infarct was measured using the Keyence Analysis software with the line measurement tool by tracing the epicardial surface of the left ventricle from base to apex for total left ventricle length and tracing from the border zone to the apex for infarct length. Infarct length was divided by left ventricular length the multiplied by 100 for percent infarct.

### Collagen quantitation

Formalin-fixed paraffin-embedded heart tissue sectioned in the coronal orientation was stained with Masson’s Trichrome following the protocol from the Center for Musculoskeletal Research at University of Rochester Medical Center. Tile scans were acquired using the Keyence BZ-x600 microscope and analyzed in ImageJ. The freehand selection tool was used to manually select the viable myocardium above the border zone in the left ventricle for the infarcted hearts. The color thresholding tool was then used to quantify the total pixel area of the viable myocardium, then the hue slider was centered between 135-180 to quantify the pixel area of collagen. The area of collagen was divided by the total area of the viable myocardium and multiplied by 100 to quantify percent area of collagen.

### Immunohistochemistry and confocal microscopy

Tissue samples were formalin fixed and paraffin embedded as previously described (see “Histological staining”).^57^ Paraffin sections (5μm) were deparaffinized, subjected to antigen retrieval in 10mM citrate pH 6.0 in TN buffer (150mM NaCl, 100mM Tris pH 7.6) for 30 minutes. Blocking was performed for 30 minutes at room temperature with 10% Horse Serum in 1xTN. Tissues were immunolabeled with antibodies Vimentin (Invitrogen cat# PA1-10003) and Cardiac Troponin I (Abcam cat# ab56357) overnight at 4°C. Fluorescently conjugated secondary antibodies were used to detect primary antibodies (donkey anti-chicken 488 Jackson Labs cat# 703-545-155 and donkey anti-goat 680 Jackson Labs cat# 705-625-147). DAPI (Thermo Fisher cat# 62248) was applied in the final wash step at 0.1μg/ml to label nuclei. Secondary antibodies were diluted 1:200. Images were acquired using a Leica SP8 confocal microscope and processed with Leica and ImageJ software. Stains were performed on at least four different samples per exposure group, and one technical replicate.

### Statistical Analysis

Data analysis and graphical representation were performed with Prism 10 (GraphPad software). Comparisons of treatment groups and sex differences was performed by two-way ANOVA. Comparisons of treatment groups with sexes pooled was performed by one-Way ANOVA. Statistical significance of interactions was identified using Tukey’s post hoc multiple comparisons test.

## Results

### Characteristics of survival and body weight following vape exposure are not altered by nicotine

The time course for vape aerosol inhalation for four weeks was intended to establish pre-existing exposure prior to myocardial infarction injury (Fig. 1A). After baseline echocardiography, an equal number of male and female mice were randomly assigned to one of three groups: 1) Control (no exposure), 2) Vape exposure, and 3) Vape with nicotine exposure. Vape aerosol exposure from Weeks 1-5 resulted in serum cotinine consistent with moderate smoking / vaping levels (Supplementary Fig. 1).^58^ After four weeks of vape exposure, an equal number of male and female mice within each of the three vape exposure groups were randomly assigned to one of two groups: 1) Sham 2) Infarct. Infarction challenge was carried out from Week 6-7 with subsequent evaluations showing no significant differences in post-infarction survival (Fig. 1C) or body weight (Fig. 1D) between the sexes or experimental exposure groups. Echocardiography was performed between Days 9 to 40 post-infarction and harvested within 24 hours.

**Figure 1.**
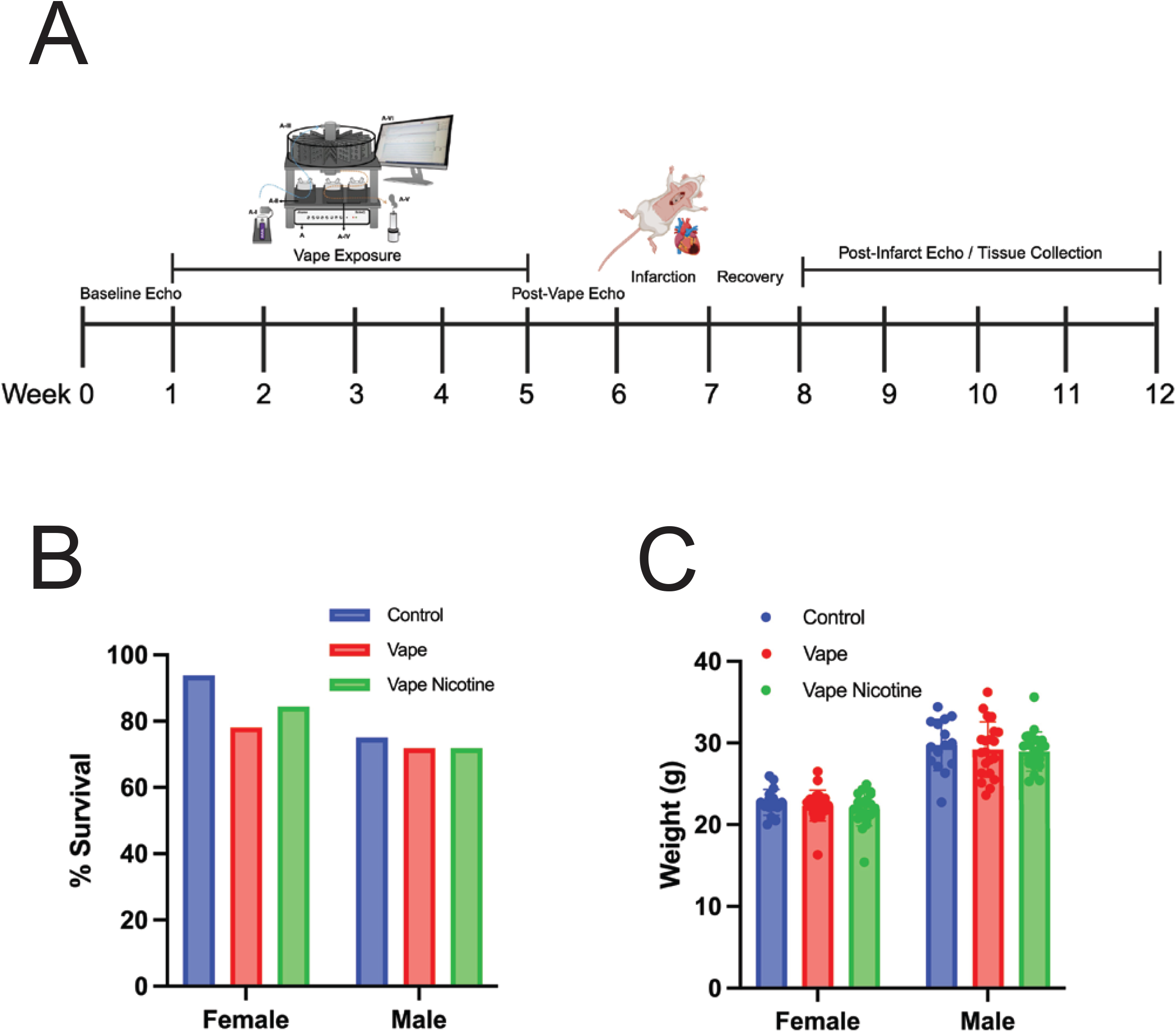
Vape exposure with or without nicotine and sex did not impact mortality or weight post myocardial infarction. A) Timeline of experiment denoting exposure, echocardiography, surgery, and sample collection. B) % of subjects that survived to the end-point of the study. C) Weight of subjects at the end-point of the study.

### Cardiac contractile performance is modestly decreased by vape exposure for one month

Three sets of echocardiographic data were acquired during the time course of the study (Fig. 2A). Baseline echocardiography was performed prior to initiation of vape exposure at Week 0-1 for initial cardiac structural and functional data. A second round of echocardiography was performed after four weeks of vape aerosol exposure to determine the impact upon myocardial structure and functional performance from Week 5-6. Myocardial infarction or sham surgery was performed on mice during Week 6-7 and a third round of echocardiography was performed between weeks 8-12 to determine the effect of vape aerosol exposure on response to the myocardial infarction. Measurements of cardiac ejection fraction were taken from long axis views at systole and diastole (Fig. 2B). Results are presented as the change in ejection fraction (EF) from baseline to post-vape (Fig. 2C,2D) or post-vape to post-infarction injury (Fig. 2E, 2F). Vape aerosol exposure did not lead to significant differences in ejection fraction between the sexes within each experimental group (Fig. 2C), so subsequent analysis was performed on the pooled male and female subjects within each group (Fig. 2D). Comparison between the experimental groups at Week 5-6 after vape exposure prior to infarction shows depressed ejection fraction in both vape aerosol groups with or without nicotine (Fig. 2D). These results are consistent with a modest but significant loss of ejection fraction resulting from vape aerosol exposure (ΔEF = -6.054 ±0.99 ; -6.199 ±0.83) for Vape or Vape Nicotine respectively versus Control (ΔEF = -3.90 ±0.79).

**Figure 2.**
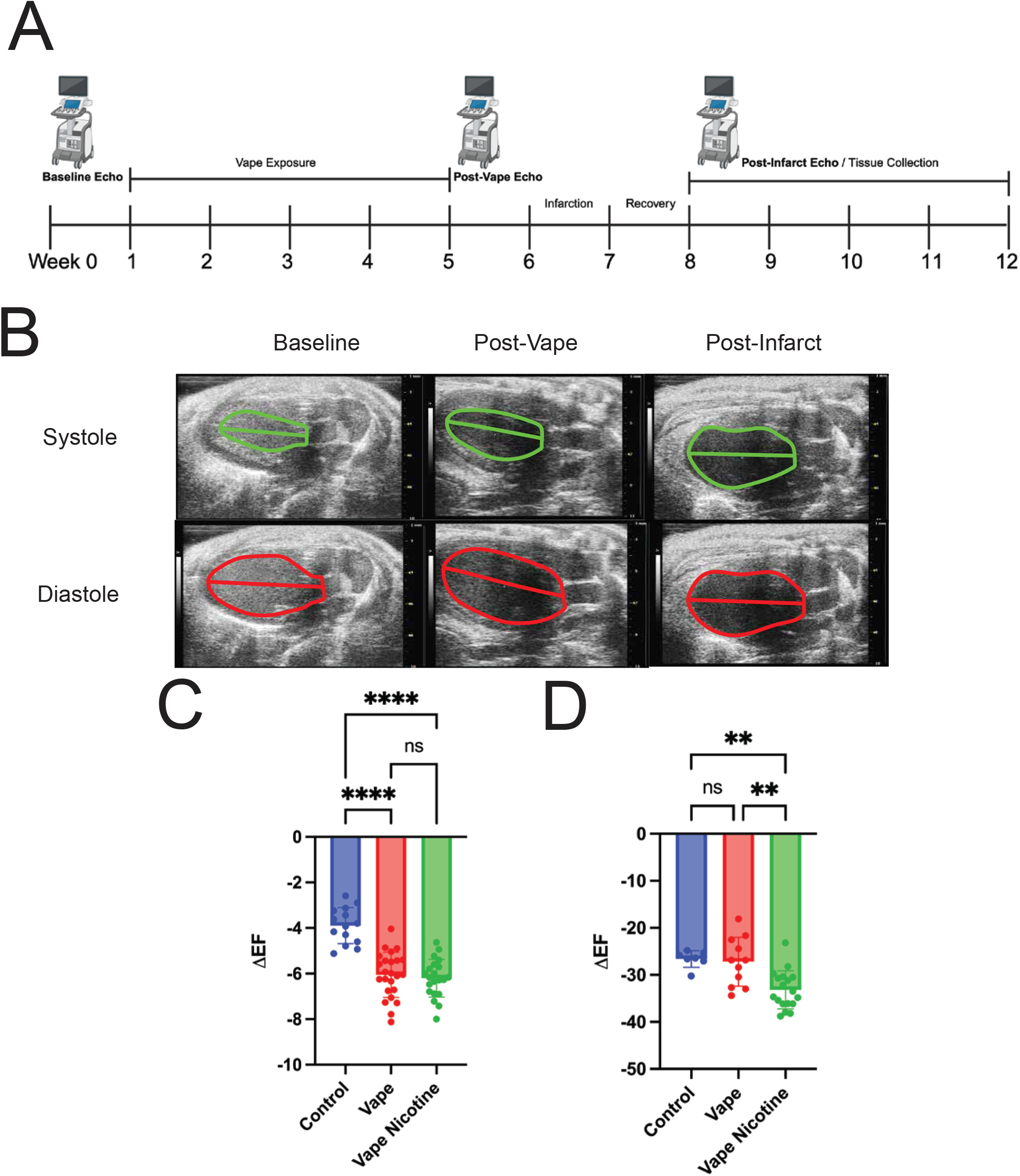
Vape exposure with nicotine impairs cardiac function post-infarction. **A)** Timeline of echocardiographic analysis. **B)** Representative images of Vape Nicotine group at baseline, post-vape, and post-infarct time points. **C)** Change in ejection fraction between baseline and post-vape time points treatment groups. **D)** Change in ejection fraction between post-vape and post-infarct time points between treatment groups. Statistical analysis = One-way ANOVA with Tukey’s multiple comparisons test (**P<0.01, ****P<0.0001)

### Cardiac contractile performance following infarction injury is worsened by vape aerosol exposure to nicotine

Following myocardial infarction injury both males and females showed comparable loss of ejection fraction (Fig. 2E), so subsequent analysis was performed on the pooled male and female subjects within each group. Comparison between experimental groups after infarction injury shows significant loss of ejection fraction in the Vape Nicotine group compared to either Control or Vape alone (Fig. 2F). In contrast, ejection fraction was comparable between Control or Vape alone groups following infarction injury. These results are consistent with a further worsening of ejection fraction output resulting from Vape Nicotine exposure (ΔEF = 33.20 ±4.05) versus Control (ΔEF = -26.62 ±1.78) or Vape alone (ΔEF = -27.21±5.20) groups.

Therefore, the presence of nicotine in the vape aerosol is responsible for exacerbating loss of myocardial function.

### Measurements of infarct size and collagen deposition are comparable between experimental groups

Hearts were processed for histopathological analyses (Fig. 1A). Infarct size measurements performed on sections stained with hematoxylin and eosin (Fig. 3A) showed comparable level of injury for all experimental groups represented as percentage of left ventricular wall involvement (Fig. 3B). Ventricular wall damage from the infarction challenge was substantial, ranging from 38.8-75.3% of the entire free wall. These findings are consistent with the profound drop in ejection fraction confirmed by echocardiography (Fig. 2) but cannot account for the observed impact of nicotine relative to the other groups. Collagen deposition measurements performed on trichrome stained sections (Fig. 3C) excluded the infarct scar region to assess whether increased fibrosis was present in the remaining viable myocardium. Collagen deposition was not significantly different between the three experimental groups. It should be noted that variability present within samples within each experimental group reflects unique pathological outcomes for individual samples as could be reasonably expected for infarction injury. Regardless of the underlying basis for this variability, the collagen quantitation also cannot account for the observed impact of nicotine on functional performance relative to the other groups.

**Figure 3.**
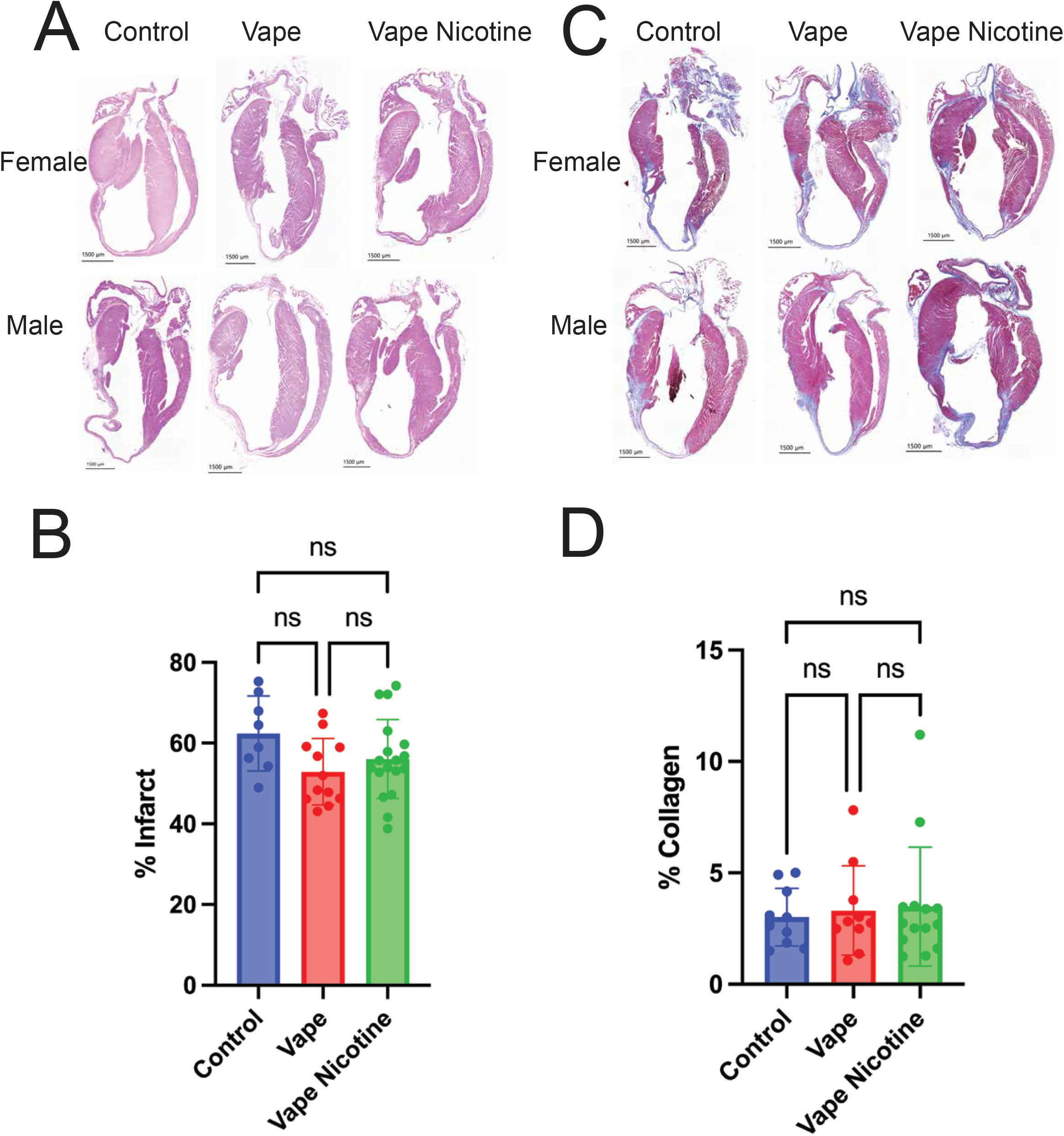
Vape exposure with or without nicotine does not affect infarct size or collagen deposition. **A)** Representative H&E stains of coronal sections of infarcted hearts. **B)** % of the left ventricular wall that is infarcted measured in H&E stains. **C)** Representative Masson’s Trichrome stain of coronal sections of infarcted hearts. **D)** % area of collagen in viable left ventricle myocardium measured in Trichrome stains. Statistical analysis = One-way ANOVA with Tukey’s multiple comparisons test.

### Cellular response to infarction injury shows differential outcomes depending upon presence of nicotine in vape aerosol

Cellular response to injury plays a critical role in post-infarction remodeling and recovery.^45^ The presence of vimentin was assessed by acquiring confocal images from three distinct regions of the infarcted heart comprised of both border zones as well as within the center of the scarred region (Fig. 4A). Vimentin immunofluorescence was quantitated in each scan and then averaged together to represent intensity signal for each individual heart section. The level of vimentin intensity signal in the Vape alone group (84.40±37.30) was significantly higher compared to either the Control (24.67+11.35; p<0.0001) or Vape with Nicotine (9.77+31.10; p<0.001). However, intensity signal was not significantly different between Control versus Vape with Nicotine groups (p>0.5). Increased vimentin intensity signal in Vape alone compared to Control is consistent with an elevated interstitial cellular response to vape aerosol exposure. In comparison, the diminished vimentin immunofluorescence intensity in the Vape with Nicotine group relative to Vape alone may be caused by nicotine-mediated inhibition of interstitial cell reactivity to vape aerosol exposure.

**Figure 4.**
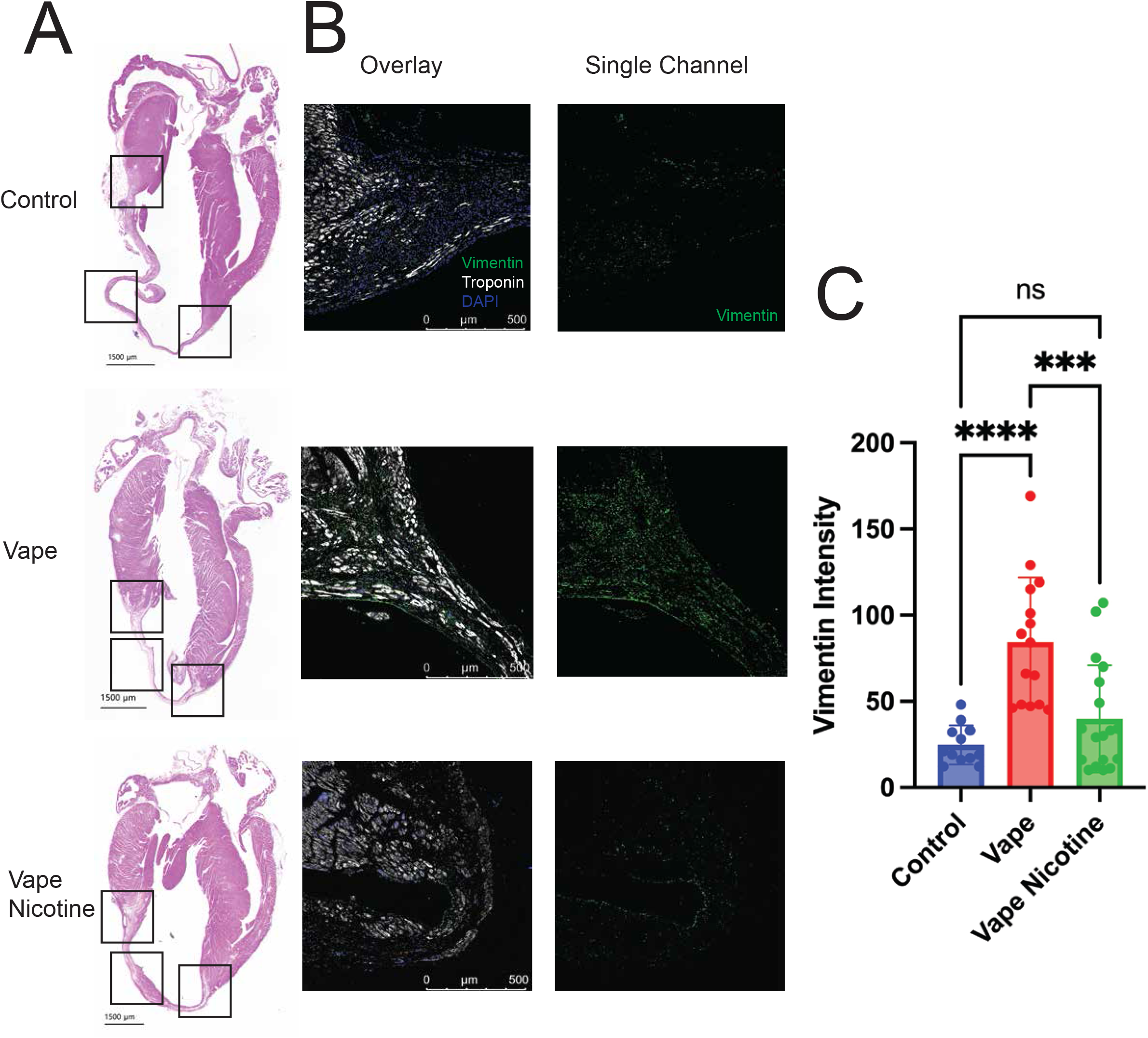
Increased vimentin expression in border zone and ischemic zone of vape-exposed infarcted hearts. **A)** Representative coronal sections of infarcted hearts (left) with boxes around high-magnification fields of view used for immunostaining and quantification. **B)** Overlay and single channel scans of border zone immunostains. White: Troponin, Green: Vimentin, Red: CD45, Blue: DAPI. **C)** Quantification of vimentin intensity in border zone and ischemic zone. 3 images per subject. N=4. Each point represents one image. Statistical analysis = One-way ANOVA with Tukey’s multiple comparisons test (*** P<0.001, **** P<0.0001).

## Discussion

The meteoric rise in vaping has been particularly distressing due to adoption by individuals with no prior history of cigarettes (the “never smokers”) as well as a youthful demographic attracted by targeted advertising and the lure of experimentation. Youthful “never smokers” lack awareness of harm caused by vaping. Furthermore, conflicting messaging from pro-vaping versus anti-vaping advocacy groups coupled with disdain or mistrust of authority figures understandably leads to indifference or skepticism regarding health-related concerns. The addictive nature of nicotine contained in cigarettes and most vape juice products leads to long term use that, rather than encouraging cessation, can escalate into greater dependency and dual use. While all parties agree that harm reduction through cessation of combustible cigarette smoking improves public health, there is abundant evidence encouraging vaping as an alternative has created a different set of problems and expanded the reach of Big Tobacco products.

Consequences of vape aerosol exposure upon myocardial structure and function present a significant health concern for e-cigarette users because of profoundly distinct biological properties of heart reparative capacity relative to the lung. The lungs are a highly regenerative organ relative to the heart, with lung tissue capable of marked recovery of function following chronic injury.^59-61^ In contrast, lack of regenerative potential in the heart results in maladaptive remodeling and deterioration of performance following damages.^62,63^ Subtle damage to the heart that may not be overtly clear in the short term can manifest decades later as loss of function leading to heart failure.^64,65^ Clinical case reports document the impact of chronic heavy vape aerosol exposure leading to cardiopulmonary damage^66^, but whether vaping predisposes the heart to impaired recovery from acute injury remains poorly understood. A recent study found that exposure to e-cigarettes during the recovery phase after infarction injury did not correlate with worsened left ventricular dilation or cardiac function^52^, a finding that contrasts sharply with similarly conceived studies assessing the effect of traditional cigarette smoking.^49^ The experimental design of these studies examine impact of exposure after myocardial infarction, but effects of cigarette smoking *prior* to myocardial injury are much more controversial. The highly debated “smokers paradox” asserts more favorable outcomes following a heart attack for cigarette smokers.^67-69^ However, to our knowledge there are no published reports evaluating the effect of vape aerosol exposure prior to acute myocardial infarction. Our findings as presented herein demonstrate vape aerosol exposure modestly impairs cardiac function either with or without nicotine in the juice in a non-injured heart, but nicotine exerts a deleterious effect upon post-infarction cardiac function as measured by ejection fraction.

Nicotine actions mediating acute physiological and biological effects upon the cardiovascular system are well known.^70^ Effects of nicotine upon myocardial recovery and healing following pathologic damage remain essentially unknown. Importantly, there is substantial cumulative literature demonstrating that nicotine impairs wound healing in other tissues including skin and bone.^71-77^ Diminished wound healing responses may result from mechanisms of nicotine action including increased vasoconstriction and decreased angiogenesis.^78^ Presumably these nicotine-mediated actions also interfere with recovery in the context of myocardial repair and recovery (and pathologic damage in general).^79^ Nicotine concentration in vape juice can produce circulating serum levels exceeding that of traditional cigarettes^54,80,81^ prompting the assertion that vaping nicotine alters myocardial healing.^79^ High concentrations and highly addictive nature of nicotine-salts used in vape juice are likely to alter myocardial repair with significant consequences for long term recovery from vaping.

Vaping can lead to deterioration of myocardial structure and function in experimental animal models^5,82^ but controversy persists regarding extrapolation of these findings to human users. Justification and / or standardization of vaping topography remains an important unresolved issue in vaping-related research. Puff parameters vary between individual reports, often without clear explanations for chosen implementation. Indeed, we carefully considered this in the design of our aerosol exposure model. Vaping topography herein is based upon published human behavior and inhalation measurements. Vaping exposure protocol is based upon experimental vaping topography recommended by Farsalinos et al. concluding “Four-second puffs with 20–30 s interpuff interval should be used when assessing electronic cigarette effects in laboratory experiments….”^21^ In addition, puff duration parameters and frequency are within the measured range of human vaping topography using electronic cigarettes in the 1 Million Puffs Study.^56^ Puff frequency, duration, and flow rate are consistent with heavy vaping consumption for humans to the extent that our chamber-based SCIREQ InExpose approach can deliver similar parameters. Validity of our exposure parameters is further reinforced by assessment of serum cotinine levels at 67.03±45.82 <u>ng/ml</u> in mice after high concentration nic-salt (50 mg/pod) vape aerosol inhalation using our protocol – a value below average for human vapers (243.72 ng/ml).^83^ Blood samples were harvested after four weeks of aerosol exposure prior to infarction for determination of cotinine serum concentration. One month of aerosol exposure was chosen as the infarction time point to avoid marked cardiopulmonary remodeling from prolonged vape aerosol exposure in our experimental model.^5^ Judicious implementation of puff topography using real-world vaping products is appropriate for this study despite limitations and compromises inherent in all animal models.

The concept of myocardial interstitial cell responses (both intrinsic and inflammatory) as determinants of the healing response following injury has gained traction in recent years.^45^ Among the numerous cardiac interstitial cell types involved in myocardial repair the role of mesenchymal cells has attracted intense interest.^84-86^ Vimentin is a cytoskeletal protein highly expressed in cells of mesenchymal origin and has been advanced as a marker of myocardial remodeling that may be valuable for diagnosis of ischemic heart disease.^87^ Furthermore, vimentin-positive mesenchymal cells enriched in the border zone of infarction were correlated with improved cardiac repair and functional performance.^88^ Similar findings for vimentin as a marker of regeneration were also reported in the kidney.^89^ Accrual of vimentin immunofluorescence in Vape alone myocardial tissue sections (Fig. 4) is unlikely to represent the acute inflammatory wave of cellular infiltrates responding to infarction injury since the hearts were harvested well after the peak inflammatory response occurring days after injury.^90-92^ Therefore, the increased vimentin intensity in the Vape alone group likely represents a persistent intrinsic mesenchymal cell response that may contribute to maintenance of ejection fraction. It is tempting to speculate that impairment of this cellular response by including nicotine in the vape juice is correlated with the worsening of cardiac function relative to Control or Vape alone groups. The connection between vape aerosol exposure and the role of nicotine in an altered myocardial cellular response post-infarction merits further investigation, as this may provide insights regarding the impact of vaping upon myocardial biology.

## Acknowledgements

MA Sussman is a recipient of funding from the California Tobacco Related Disease Research Program (Pilot award T31IP1790 and Impact award T31IR1585). The author extends his deepest appreciation to members of the Sussman Laboratory who provide invaluable assistance and expertise in developing vaping-related studies and information.

## Abbreviations

ENDS: Electronic nicotine delivery system
PG: Propylene glycol
VG: Vegetable glycerin

## Notes

### Competing Interest Statement

The authors have declared no competing interest.

