## Supplementary figures and images for "Myocardial infarction injury is exacerbated by nicotine in vape aerosol exposure"

### Supplemental Figure 1

Supplementary Figure 1

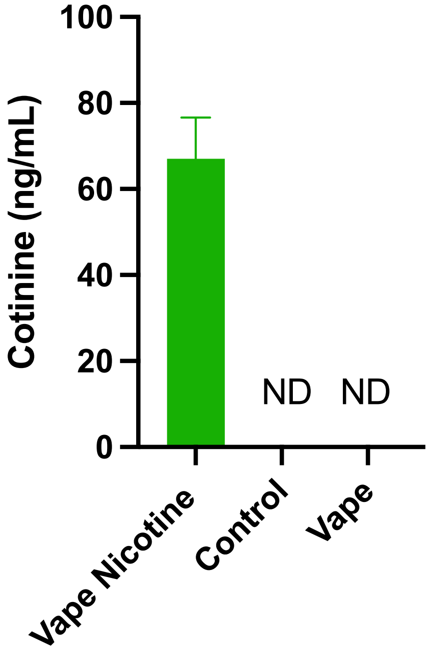
